# Turnip mosaic virus drives selective filtering and community reassembly in the *Arabidopsis thaliana* root microbiome in a genotype-specific manner

**DOI:** 10.64898/2025.12.11.693707

**Authors:** Alberto Cobos, Zulema Udaondo, Irene Gonzalo, Gabriel Castrillo, Adrián A. Valli

## Abstract

Plant root microbiomes play a central role in plant health, yet their responses to viral infection remain poorly understood. Here, we investigated how Turnip mosaic virus (TuMV) alters root-associated bacterial and fungal communities in two *Arabidopsis thaliana* genotypes (Col-0 and Mar-12) grown in natural soils. Using 16S and ITS amplicon sequencing, we assessed changes in diversity, taxonomic composition, enriched microbial taxa, and co-occurrence network structure to distinguish between plant-mediated recruitment (“cry-for-help”) and pathogen-induced dysbiosis.

TuMV infection caused a pronounced reduction in bacterial diversity and a restructuring of bacterial community composition, whereas fungal communities remained largely stable. Viral infection also led to genotype-specific shifts in enriched bacterial genera, with opportunistic and stress-tolerant taxa proliferating differently in each genotype. Despite the initial perturbation, bacterial networks recovered connectivity and, in some cases, reached higher complexity than those of healthy plants, indicating strong microbial resilience.

Together, these results reveal that TuMV infection acts as a selective filter on bacterial, but not fungal, root communities and that the surviving taxa can reorganize into functional networks. Our study provides one of the most comprehensive assessments of virus-induced microbiome restructuring and highlights the importance of host genotype in shaping microbial responses to biotic stress.

## 1. Introduction

The plant root microbiome is a critical determinant of plant health, influencing nutrient acquisition, growth, and resistance to stress through a complex network of interactions among microbes, the host and the soil environment. Roots are not passive surfaces: they exude a chemically diverse cocktail of low- and high-molecular-weight compounds that shape the surrounding microbial community by providing resources, signalling molecules and selective pressures that favour particular taxa and functions (Feng et al., 2024; Robert et al., 2025). Root-associated microbes in turn affect plant physiology through nutrient cycling, hormone modulation and antagonism against pathogens, making the rhizosphere and endosphere loci of dynamic plant–microbe co-evolution (Ali et al., 2023; Trivedi et al., 2020). Because this interaction network is both functionally important and ecologically intricate, understanding how microbial communities assemble around roots has become a central focus of plant microbiome research. Community composition and assembly reflects the balance of deterministic forces (microorganism selection by the plant root exudates) and stochastic processes (ecological drifts or niche rearrangements), and the relative importance of these factors can shift with context such as development stage, soil type or environmental stress (Hayashi et al., 2024).

Stress, both abiotic and biotic, frequently remodels root microbiomes. Drought, nutrient limitation, soil contamination and herbivore/pathogen attacks have all been shown to alter diversity, taxonomic composition and network structure of root-associated communities, sometimes in ways that impair plant function and sometimes in ways that increase disease suppression or tolerance (Robert et al., 2025). Regarding biotic stresses, there are mainly two divergent outcomes over microbe populations, a *directional plant-mediated recruitment* (the so-called “cry-for-help”) and *non-directional plant-independent restructuring* (pathogen induced dysbiosis). The cry-for-help idea proposes that stressed plants modify root exudation and other signals to selectively recruit microbes that can assist in defence or stress mitigation, thereby assembling a protective rhizobiome; this hypothesis is supported by mechanistic work documenting stress-induced changes in exudate chemistry and by field and microcosm studies that find enrichment of taxa with putative beneficial traits after attack (Rizaludin et al., 2021; Rolfe et al., 2019). In contrast, dysbiosis refers to an unstructured loss of community stability or function, often marked by reduced diversity and weakened network complexity, that arises when biological stress overwhelms host control or perturbs ecological interactions. Understanding these processes could be crucial to unravel the factors and interactions leading to the selection of beneficial microorganisms or the absence of it (Liu et al., 2023; Rizaludin et al., 2021).

As described above, biotic stresses can drive root microbiomes along markedly divergent trajectories. On one hand, several clear cases of dysbiosis have been documented. For example, a recent study in common bean infected by *Fusarium oxysporum* showed that infection significantly altered both rhizosphere and endosphere microbiota, restructuring community composition and function in a cultivar-dependent manner (Mendes et al., 2023). Another study on bacterial wilt caused by *Ralstonia solanacearum* found that diseased plants exhibited reduced fungal diversity in rhizoplane and endosphere, together with increases in pathotrophic fungi and losses of key community members (Tao et al., 2025). Similarly, soybean plants suffering from root disease showed a decline in fungal diversity and an expansion of opportunistic taxa in their root-associated microbiota (van Bentum et al., 2025). On the other hand, several studies support the idea that pathogen invasion can trigger protective microbial recruitment. In a wheat - *Rhizoctonia solani* system, repeated cultivation under pathogen pressure led to the accumulation of pathogen-antagonistic bacterial taxa, which were associated with lower disease incidence (Yin et al., 2021). In another report, infection of sugar beet roots induced enrichment of potentially beneficial microbes in the rhizosphere, and a reconstituted community from diseased plants suppressed disease in controlled experiments (Gao et al., 2021). Moreover, in banana seedlings grown in soils with a history of *Fusarium* wilt, endophytic *Pseudomonas* strains reduced pathogen survival and promoted beneficial microbial groups (Lv et al., 2023). Together, these contrasting examples show that pathogen invasion can lead either to destabilizing microbiome compositions or to reassembling toward protective communities, depending on soil context, host background and the microbial taxa available for recruitment.

Mechanistically, this directed recruitment can be explained by the intimate cross talk between plant and microbiota via root exudates. Root exudation responds plastically to environmental cues, and the chemical shifts can include carbohydrates, amino acids, phenolics and secondary metabolites that act as chemoattractants, nutritional subsidies or antimicrobial, thereby reshaping the pool of microbes able to colonize the rhizosphere and the root surface. These exudate-mediated filters, together with plant immune modulation and microbe-microbe interactions, create routes by which plants can bias assembly toward protective functions under stress (Afridi et al., 2024; Robert et al., 2025). In such cases, beneficial bacteria and fungi can trigger systemic plant immune responses in a process known as induced systemic resistance that prime the plant to future infections (Pršić & Ongena, 2020). Microbiota can also provide resistance in a direct manner by competing and inhibiting the growth of pathogens, for example, via the production of antibiotics, fungal cell wall-degrading enzymes, and siderophores (Pieterse et al., 2014; Vannier et al., 2019). At the same time, host genotype and soil origin are powerful modulators of microbiome composition and the trajectory of assembly following perturbation. Different plant genotypes may vary in their exudation profiles, immune responses and root architecture, and these differences can lead to genotype-dependent recruitment patterns and disease outcomes. Likewise, the bulk soil microbial pool and soil physicochemical context determine which taxa are available to be recruited and how interactions play out. Consequently, experiments that cross host genotypes with distinct soil inoculums are particularly informative for teasing apart host-driven selection from source-pool constraints (Feng et al., 2024; Z. Wang & Song, 2022).

Despite the growing body of work on bacteria and fungi, studies that explicitly examine how viral infection alters root microbiomes remain relatively scarce. Viruses are historically under-represented in plant research (Rubio et al., 2020), partly because their intracellular lifestyle and detection challenges made them less obvious candidates for direct microbiome studies, but recent amplicon-based and network analyses indicate that viral infection can cascade to reshape root-associated bacterial communities, reduce diversity and simplify co-occurrence networks, as shown for several crop systems. The comparative scarcity of virus-focused microbiome studies means that general patterns and mechanisms are still poorly resolved, and that systematic experiments are needed to assess whether virus-driven changes are more often directional (Li et al., 2025) or symptomatic of dysbiosis (Y. Deng et al., 2024; Liu et al., 2023; F. Wang et al., 2024; N. Wu et al., 2024).

Taken together, the literature indicates three points that motivate the present work. First, root microbiomes are central to plant health and are shaped by an interplay of plant physiology, soil context and microbial interactions (Feng et al., 2024; Robert et al., 2025). Second, biotic stress can reconfigure these communities in ways that are functionally meaningful, but whether those changes constitute adaptive recruitment (cry-for-help) or non-adaptive dysbiosis remains an open empirical question that requires joint taxonomic, functional and network analyses (Afridi et al., 2024; Rizaludin et al., 2021). Third, virus-induced microbiome changes are understudied, and controlled microcosm experiments that use inoculum from natural soils offer a pragmatic route to examine causality while retaining ecological realism (Liu et al., 2023; Mohamed et al., 2024). In this study, we challenged two *Arabidopsis thaliana* genotypes against TuMV infection, exposing them with microbiota coming from a location where Arabidopsis naturally grow. We examined the epi-endosphere, profiling bacterial and fungal communities through 16S and ITS sequencing. Our approach integrated analyses of alpha and beta diversity, enrichment of taxa and functions, and co-occurrence network structure to determine how viral infection reshapes microbial communities. We hypothesize that TuMV alters the root microbiota and tested whether such changes reflect a plant-mediated cry-for-help response or a plant-independent restructuring associated with virus induced dysbiosis. We also examined whether plant genotype modulate these microbiome shifts. By combining multiple genotypes, natural soils, and studying bacterial and fungal communities, our experimental design provides higher ecological resolution than many previous studies and establishes a robust foundation for understanding the ecological implications of dysbiosis and the potential for microbiome-mediated mitigation of viral disease in natural and agricultural systems.

## 2. Material & methods

### 2.1 Soil, plant and virus material

The soil field samples were collected from three locations in central Spain: Pinar de Valdelatas, Madrid (VAL, 40°32’12.62”N, 3°41’29.26”O); Ciruelos de Coca, Segovia (CDC, 41°12’48.73”N, 4°32’44.33”O) and Marjaliza, Toledo (MAR, 39°34’52.79”N, 3°55’53.90”O). *Arabidopsis thaliana* genotypes Col-0 (Columbia, USA) and Mar-12 (Marjaliza, Spain) were used. An infectious clone of the JPN1 strain of TuMV (pGTuMVJPN1-GFP, DOI: 10.1007/s10658-016-1057-9) was propagated in *Nicotiana benthamiana* plants and crude extracts from infected leaf tissues were used for further inoculation.

Microcosm soils were set up by inoculating sterile greenhouse substrate (peat:vermiculite, 3:1) with bulk soil liquid extract. These extracts were prepared by diluting the soil in PBS buffer, ensuring a volume to reach a minimum of 10^6^ CFU per mL of greenhouse substrate. Each bulk soil CFUs were estimated prior to the inoculations by plate serial dilutions. In parallel, equivalent volumes of the extracts were sterilized to stablish the Mock soil treatments. Plants were grown in controlled chambers at 21ºC with short day conditions (8h/16h, light/dark) for two months and with long day conditions (16h/8h, light/dark) for one month, to allow Mar-12 plants flowering.

### 2.2 Experimental design

The first step was the soil inoculation, each soil type having a Mock soil treatment, resulting in 6 soil treatments (VAL, VAL-Mock, CDC, CDC-Mock, TOL & TOL-Mock). Then, 24 plants per genotype were transplanted to individual pots per soil treatment (12 in the case of the Mock soils). Finally, half of the plants from each soil type were mechanically inoculated with TuMV. At the end of the experiment, root and leaf samples were collected for DNA extraction and western blot viral detection respectively.

### 2.3 Virus inoculations

TuMV mechanical inoculations were carried out using N. *benthamiana* infected leaves ground in 5mM pH 7.2 phosphate buffer in a 1:4 proportion (w:v) with an ice-cold mortar and pestle, and the sap was finger-rubbed onto three leaves that had previously been dusted with Carborundum. Mock plants were inoculated with just the phosphate buffer. All inoculations were performed at Arabidopsis developmental stages 1.06 (Boyes et al., 2001). The efficiency of inoculations was determined by detecting systemic virus GFP signal in all plants.

### 2.4 Immunodetection of proteins by western blot

The preparation of protein samples under denaturing conditions, the separation on SDS-PAGE and the electroblotting to nitrocellulose membranes was done following the protocol as recommended by the manufacturers (iBlot3, Invitrogen). TuMV was detected using anti-CP (Ref. AS-0132, DSMZ) as primary antibody and horseradish peroxidase (HRP)-conjugated goat anti-rabbit IgG (Ref. 31466, Thermo Fisher Scientific) as the secondary reagent. Immunostained proteins were visualized by enhanced chemiluminescence detection with Clarity ECL Western blotting substrate (Bio-Rad) in a ChemiDoc apparatus (Bio-Rad). Ponceau red was used to verify equivalent loading of total proteins in each sample.

### 2.5 Root DNA extraction and sequencing of the 16S rRNA gene

Root surface was thoroughly cleaned of any solid residue with running water and then washed three times with sterile PBS buffer, keeping just the epi/endophytic microorganisms. The clean roots were grounded in liquid nitrogen and stored at −80°C for further treatment.

According to the manual, root DNA was extracted using the DNeasy PowerSoil Pro kit (Qiagen, Germany). The quality of the DNA extracts was evaluated with a NanoDrop Spectrophotometer (NanoDrop Technologies, Wilmington, DE, USA). Bacterial 16S rRNA gene fragments (V5–V7) were amplified from the extracted DNA using primer sets 799F (5⍰-AAC MGG ATT AGA TAC CCKG-3⍰) and 1193R (5⍰-ACG TCA TCC CCA CCT TCC-3⍰) as these primers should amplify less host DNA than other pairs (REFERENCE). The DNA amplification and sequencing were done using an Illumina NovaSeq X Plus by Novogene (Beijing, China).

### 2.6 Sequence processing of 16S rRNA gene

Paired-end reads were quality-checked using QIIME 2 (Bolyen et al., 2019) demux summarize visualization. Quality profiles showed consistently high Phred scores across all positions (median ≈ Q37, lower quartile ≥ Q25), with no appreciable drop in quality at read ends. Given the uniformly high read quality, no trimming or truncation was applied (trim-left = 0, trunc-len = 0). Denoising, paired-end merging, and chimera removal were performed with the DADA2 plugin using default parameters, including consensus-based chimera detection. DADA2’s error-model learning and quality filtering removed only the small fraction of low-quality reads (≈2%), yielding high-confidence amplicon sequence variants (ASVs) for downstream analyses.

Amplicon sequence variants (ASVs) were taxonomically classified using the QIIME 2 feature-classifier plugin with a naïve Bayes classifier. Bacterial 16S rRNA sequences were annotated against the SILVA 138 database, whereas fungal ITS sequences against the UNITE v10 dynamic database. Finally, ASVs classified as mitochondria or chloroplast were removed for further analysis.

### 2.7 Statistical analysis

#### 2.7.1. Diversity analysis

Alpha-diversity indexes including Shannon index, Pielou’s evenness index and Faith’s index were calculated using the normalized ASV table in QIIME2 and then visualized in box plots. Kruskal-Wallis rank-sum test included in the QIIME2 package was used to compare the significant difference between alpha indexes of bacterial and fungal communities in healthy and diseased samples.

Beta diversity was analysed using Bray-Curtis dissimilarity to assess compositional differences between samples. Resulting distance matrices were visualized via Principal Coordinates Analysis (PCoA) to provide an intuitive graphical representation of sample similarity. Statistical significance of community separation was evaluated using PERMANOVA included in the QIIME2 package, providing quantitative evidence of differences between groups.

#### 2.7.2. Taxon enrichment analysis

Differential abundance analysis of ASVs were performed using multiple complementary statistical approaches to ensure robustness of the results. Low abundance features were first filtered to remove rare ASVs that could bias the analyses. Differential abundance was then evaluated using three independent methods. ANCOM-BC implemented from the QIIME2 package, DESeq2 (Love et al., 2014) and ALDEx2 (Fernandes et al., 2013) R packages. Each method is based on different statistical principles: compositional bias correction, negative binomial modelling, and centered-log ratio transformation with Monte Carlo sampling, respectively. Only ASVs identified as significantly enriched by at least two of the three methods were considered robustly differentially abundant. Bar plots were made with ggPlot R package (Wickham, 2009).

#### 2.7.3. Functional enrichment analysis

Functional enrichment analysis was performed using PICRUSt2 to infer metagenome content from the ASVs tables and to predict KEGG Orthologs (KOs) and metabolic pathways. Differential abundance of predicted functions between conditions was evaluated using both parametric (t-test) and non-parametric (Mann–Whitney U) comparisons, retaining KOs that were significant under at least one test (p < 0.05). Significant KOs were annotated using the KEGG REST API, and associated pathways were identified through KO–pathway mapping files. Pathway-level differences were additionally assessed using hypergeometric enrichment tests, with FDR correction applied to control for multiple testing. Finally, enriched genes and pathways were visualized using dot plots and heatmaps to highlight functional shifts between conditions.

### 2.9 Co-occurrence network construction

Bacterial co-occurrence networks were constructed using the Molecular Ecological Network Analyses Pipeline (Y. Deng et al., 2012). Briefly, ASVs with an average relative abundance >0.05% and occurring in at least 4 replicates were retained for network inference. The network construction was based on recommended parameters in data preparation and random matrix theory default settings. Bacterial co-occurrence networks were visualized with Cytoscape (v3.10.4) (Shannon et al., 2003). The network modules were characterized by greedy modularity optimization. Keystone species were identified based on the values of within-module connectivity (Zi) and among-module connectivity (Pi) according to a previous study (Y. Deng et al., 2012). Nodes (ASVs) could be classified into four categories, including peripheral nodes (Zi⍰≤⍰12.5 and Pi⍰≤ ⍰0.62), connectors (Zi⍰≤ ⍰2.5 and Pi⍰> ⍰0.62), module hubs (Zi⍰> ⍰2.5 and Pi⍰≤ ⍰0.62), and network hubs (Zi⍰> ⍰2.5 and Pi⍰> ⍰0.62).

## 3. Results

### 3.1 General characterization of the endophytic dataset

To characterize the microbial amplicon sequence variants (ASVs) within *Arabidopsis thaliana* roots, bacterial (V5–V7 16S rRNA) and fungal (ITS1–1F) regions were sequenced from epi- and endophytic layers of roots grown in three distinct soils and coming from two host genotypes.

After quality filtering and OTU clustering, rarefaction curves approached saturation for all samples, confirming sufficient sequencing depth for both bacterial and fungal samples (Figure S1).

The taxonomic composition was well resolved at the genus level, with consistent representation across higher taxonomic ranks, ensuring robust comparisons across soils, host genotypes and infection stress (Figures 1A & 2A, for bacteria and fungi respectively).

**Figure 1.**
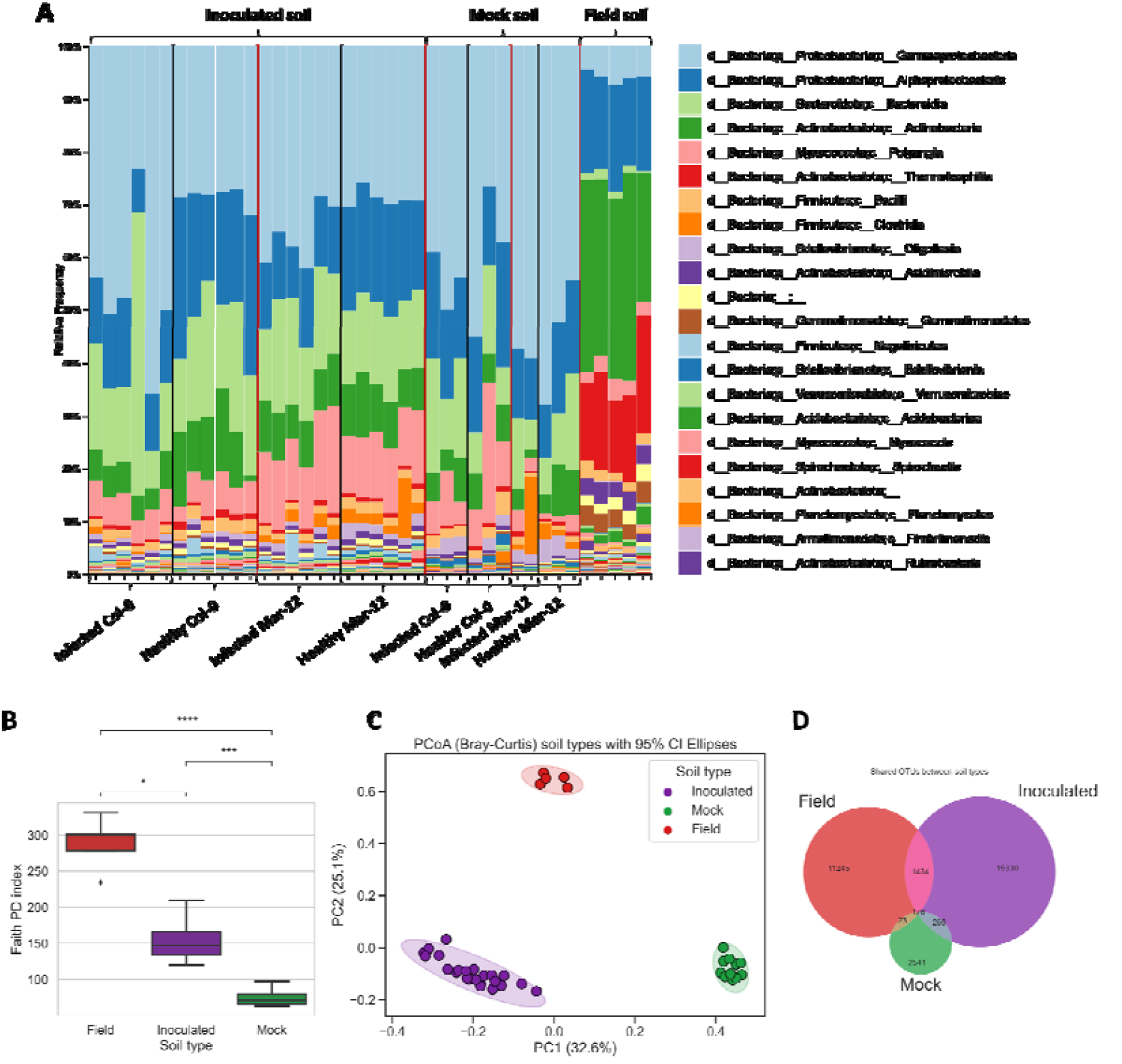
General description of the bacterial communities regarding their soil type. A) Histogram depicting the proportional presence of bacterial ASVs. B) Alpha diversity observed in each soil type. C) PCoA representing the community structuring based on soil type. D) Venn diagram showing the shared ASVs between soil types.

**Figure 2.**
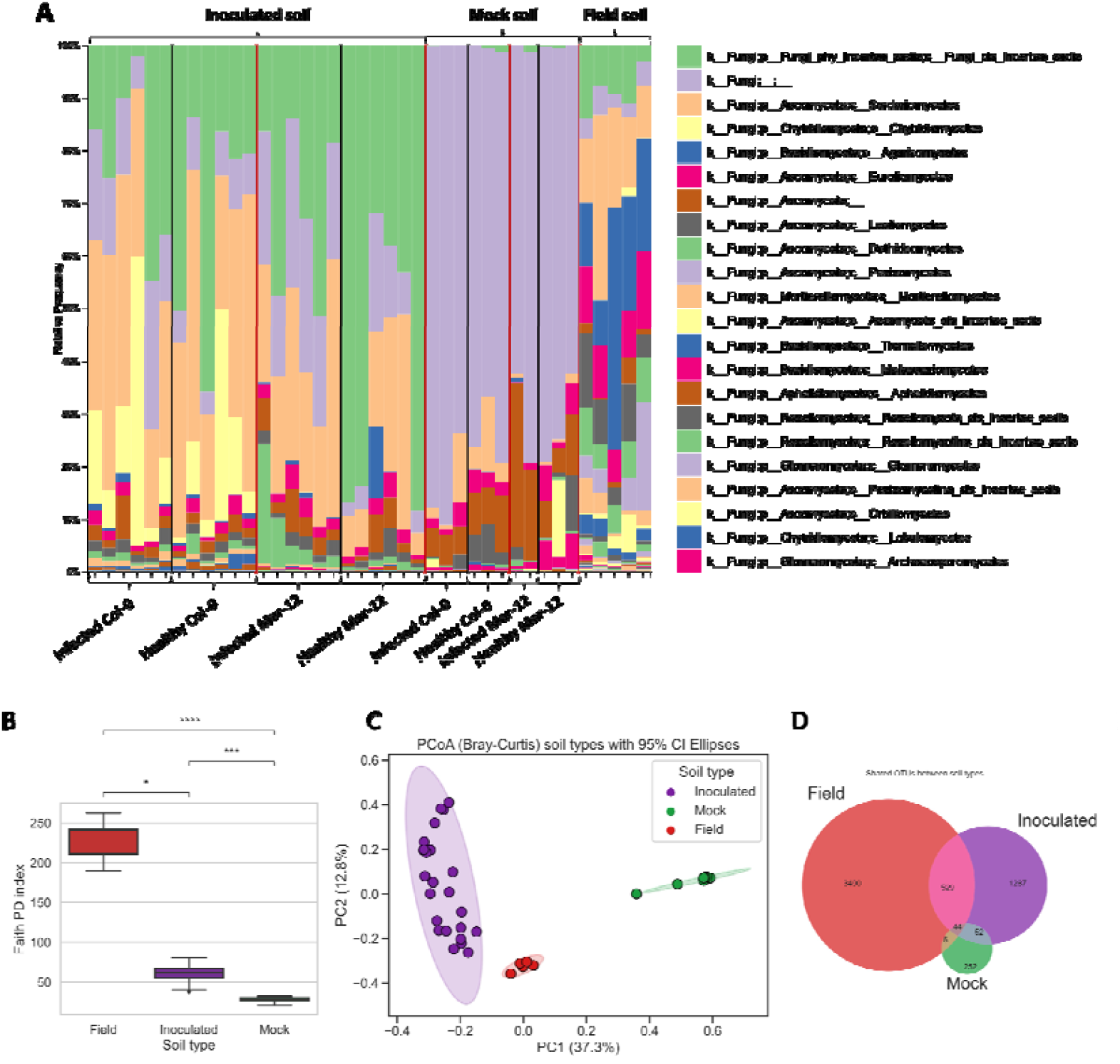
General description of the fungal communities regarding their soil type. A) Histogram depicting the proportional presence of bacterial ASVs. B) Alpha diversity observed in each soil type. C) PCoA representing the community structuring based on soil type. D) Venn diagram showing the shared ASVs between soil types.

Prior to conducting further analysis, we checked the quality of our microcosm soils. The three types of inoculated soils were well differentiated (Figure 3: PCoA). Bulk soil field samples were the most diverse, followed by the inoculated soils (Figures 1B & 2B, for bacteria and fungi respectively). Groups were well differentiated as shown with the PCoAs and PERMANOVA (Figures 1C & 2C, for bacteria and fungi respectively). Finally, the number of shared ASVs was significantly higher between bulk and inoculated soils than between inoculated and mock soils (Figures 1D & 2D, for bacteria and fungi respectively).

**Figure 3.**
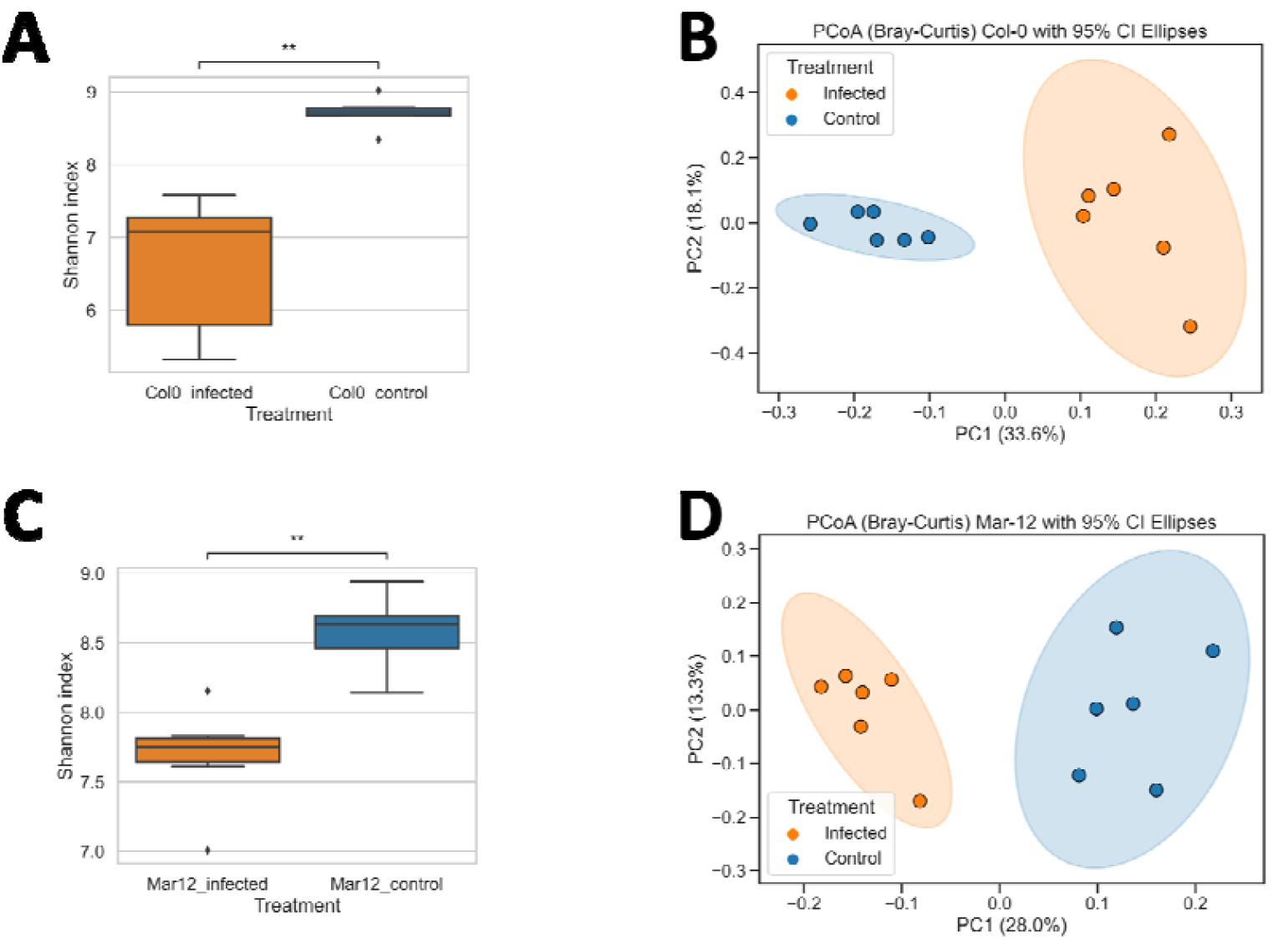
Genetic diversity and population structuring of bacterial ASVs. A) Alpha diversity comparing infected with healthy Col-0 plants. B) comparing infected with healthy Col-0 plants C) Alpha diversity comparing infected with healthy Mar-12 plants. D) comparing infected with healthy Mar-12 plants.

This suggests that microorganisms coming from the field were more impactful in the assembly of the observed communities than the ones coming from the greenhouse environment or irrigation.

### 3.2 Genotype-dependent modulation of microbial communities

As described in previous studies, plant genotype or cultivar could be an important factor in the recruitment of root microbiota so we studied this effect with our two Arabidopsis genotypes.

In both bacterial and fungal communities, plant genotype didn’t affect the biodiversity as reflected by alpha diversity indexes (Figure X1). On the other hand, beta diversity distances showed significant differences in the community composition between genotypes (Figure X2). This trend was observed in all plants regardless of the presence of the virus. Moreover, differential abundance analysis confirmed that taxa enrichment was genotype specific as virus infection enriched taxa in Col-0 showed no effect in Mar-12 and vice versa (Figure X3).

These findings indicate that host genotype modulates the extent and nature of microbiome recomposition independently of the presence of stress. Moreover, the initial presence of a minimal microbial community appears to be needed for this modulation to occur, as no differences were found between genotypes when studying healthy plants growing on Mock soils (Data not shown). Based on these observations, all following analysis will be carried out for each genotype independently.

### 3.3 Viral infection alters endophytic microbial diversity across soils and genotypes

#### 3.3.1 Diversity analysis

Alpha- and beta-diversity analyses revealed significant effects of viral infection on the structure of the endophytic microbiome.

Alpha diversity indices for bacterial communities tended to be higher in roots from healthy plants than from TuMV-infected plants (Figure 3A & 3C). In the case of fungal communities however, the effect of TuMV infection was milder, resulting in significant differences in alpha diversity only for Mar-12 plants (Figure 4A & 4C). This reduction in biodiversity upon infection was generally significant in both host genotypes, although the magnitude varied depending on soil origin, being clearer in the case of Cat soil.

**Figure 4.**
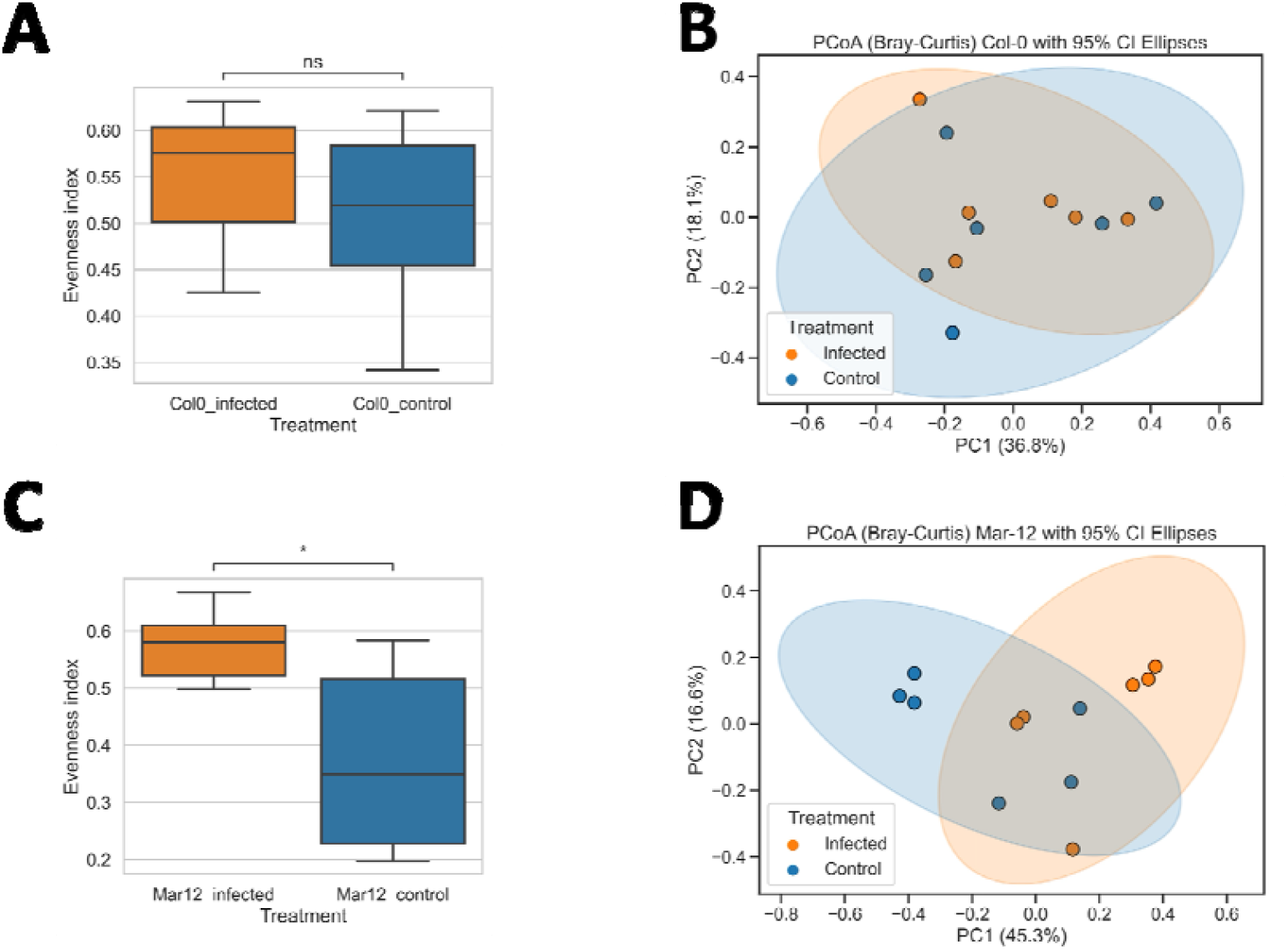
Genetic diversity and population structuring of fungal ASVs. A) Alpha diversity comparing infected with healthy Col-0 plants. B) comparing infected with healthy Col-0 plants C) Alpha diversity comparing infected with healthy Mar-12 plants. D) comparing infected with healthy Mar-12 plants.

Beta diversity analysis showed that the bacterial community structure was significantly altered by the infection. Within each soil type, infected samples formed distinct clusters from healthy controls, indicating compositional shifts associated with viral invasion (Figure 3B & 3D). On the other hand, fungal community structures were unaffected by the viral infection (Figure 4A & 4C).

These results indicate that while the local edaphic context shapes the baseline structure of the endophytic microbiome, viral infection introduces a secondary but consistent source of compositional divergence over bacterial communities.

#### 3.3.2 Differential enrichment of taxa reveals targeted compositional shifts under viral infection

Comparative analyses at the genus level identified several taxa whose relative abundance was significantly altered in response to TuMV infection. Similar to the previous results, viral infection had a stronger effect over bacterial populations. As stated in the methods section, ASVs were only considered as significantly enriched if they were detected as such by at least two of the three applied methods.

In the case of Val soil Bacteria genera such as *Actinotalea, Pelosinus* and *Rhizobacter* were enriched in Col-0 infected plant roots, whereas *Cytophaga, Iamia* and *Paenibacilllus* were depleted when compared to healthy roots. In addition, genera like *Methylotenera, Flavobacterium* and *Pajaroellobacter* increased in virus infected Mar-12 roots, while *Rhodopseudomonas* and *env*.*OPS 17* were less abundant in these plants. Finally, it is important to highlight that the genera *Haliangium* was enriched in both plant genotypes during infection with the same 4 ASVs supporting this effect. Regarding enriched fungi, no significant results were obtained for Col-0, while *Malassezia restricta* and *Acremonium sclerotigenum* were more abundant in infected Mar-12 roots.

Regarding Tol soil, we only found significant bacterial depletion in infected plants but no enrichment (Figure 14: enrichment barplot). The genera *Arthrobacter* and family *Oxalobacteraceae* were less present in TuMV infected Col-0 plants and the genera *Paludibacter* in the case of Mar-12. Focusing on fungi, we observed the same pattern, with the species *Neopyrenochaeta acicula* and *Fusarium nygamai* being significantly less represented in infected Col-0 roots (Figure 15: enrichment barplot).

#### 3.3.3 Microbial functional following viral infection

To assess whether compositional changes were accompanied by functional rearrangements, metagenome inference using PICRUSt was performed.

The predicted functional profiles displayed clear separation between infected and control samples in PCoA space, particularly within the bacterial dataset.

Pathways related to stress response, reactive oxygen species detoxification, and carbohydrate metabolism were significantly enriched in TuMV-infected plants, whereas functions associated with nutrient exchange (e.g., nitrogen fixation, siderophore biosynthesis) were reduced.

Overall, these results indicate that TuMV infection not only alters the taxonomic composition of the endophytic microbiome but also reshapes its predicted metabolic landscape, consistent with a community-level shift in ecological strategy.

### 3.4 Microbial network dynamics following viral infection

Four bacterial co-occurrence networks were constructed among bacterial taxa at the ASV level, plant genotype and infection combination. In parallel, other four networks were made for the fungi ASVs following the same criteria.

Constructed networks did not differ much in terms of complexity and cohesion between healthy and infected roots (Figures 5 & 6). Both bacterial and fungi networks showed similar node degree, closeness centrality and betweenness centrality (Figure 6). Modularity analysis followed the same trend, with no significant effects of the infection on the Zi-Pi classification of the nodes (Figures 5 & 6). Nonetheless, bacterial networks allowed for a better classification of their nodes in terms of modularity, suggesting that infected plant networks were slightly more complex in terms of node connectivity.

**Figure 5.**
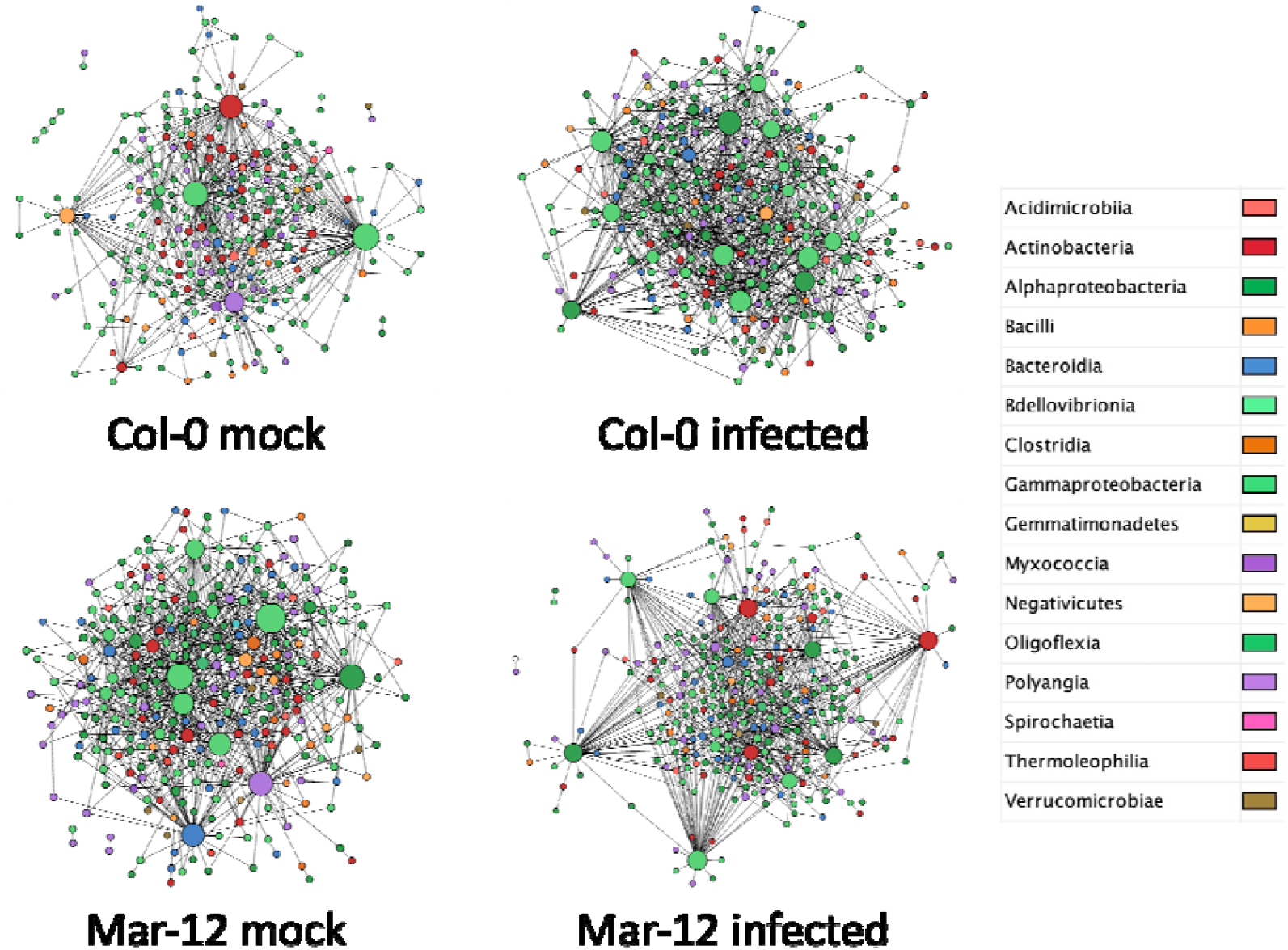
Co-occurrence networks constructed with bacterial ASVs. Colour legend represents the bacterial class.

**Figure 6.**
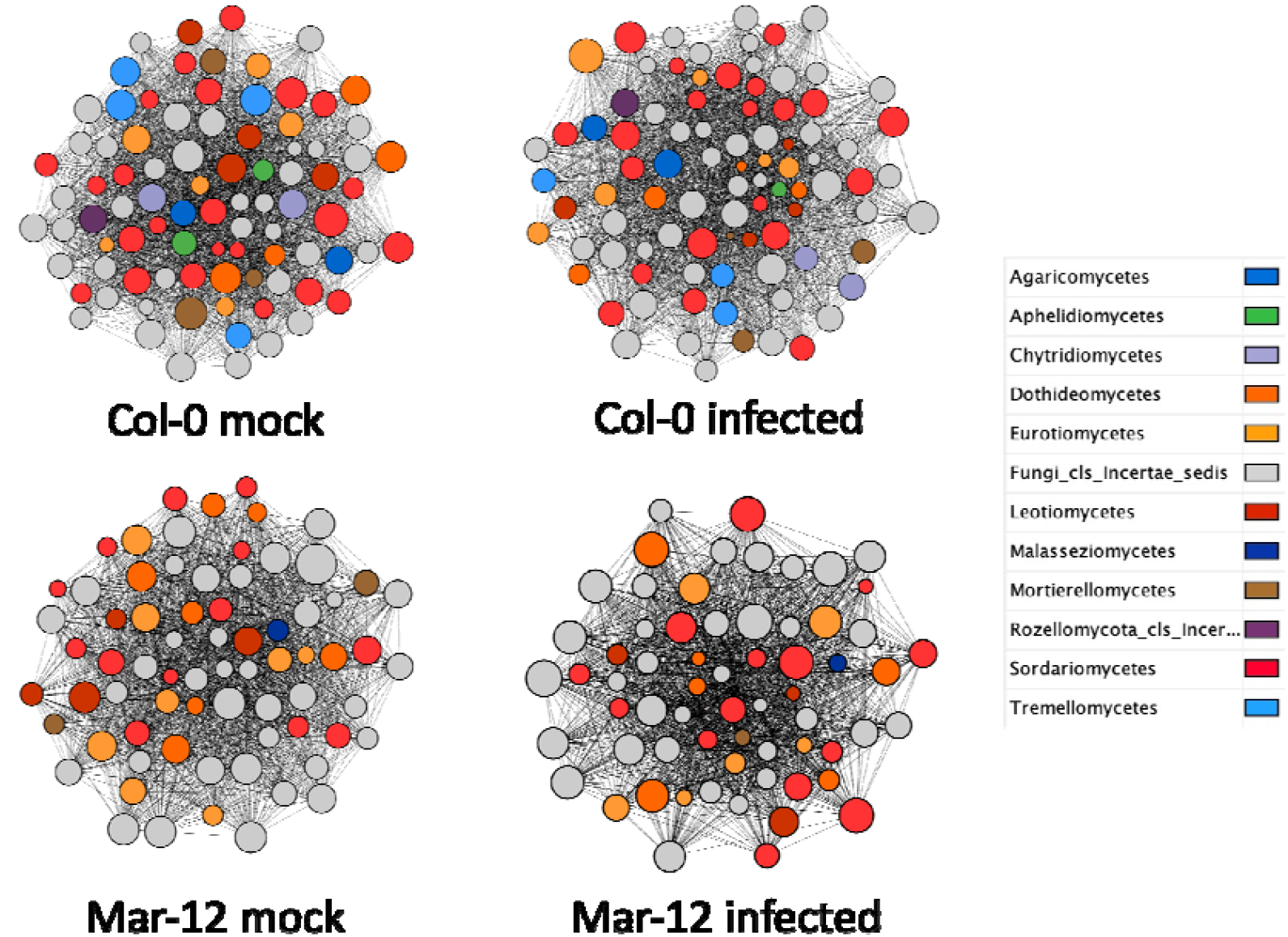
Co-occurrence networks constructed with fungal ASVs. Colour legend represents the fungi class.

Include the statistical tests comparing general node betweenness and closeness centrality (possibly without figure).

Compare the modularity of the healthy vs. infected networks (Figure representing all the nodes across the four categories coloured by treatment).

The bacterial classes involved in the Col-0 network interaction were mainly *Alphaproteobacteria* and *Gammaproteobacteria* being around 32% and 29%, respectively. The same classes were the most abundant in the case of Mar-12 with a 27% and 28% of contribution respectively. On the other hand, the most prevalent classes in the fungi networks were *Sordariomycetes* and *Eurotiomycetes*, with a 25% and 7% presence respectively in the case of Col-0. Both classes were also the most abundant in the case of Mar-12, comprising about 19% and 11% respectively.

On the other hand, the main fungal classes involved in the Col-0 and Mar-12 networks were *Sordariomycetes* and *Eurotiomycetes* being around 19-25% and 7-11% respectively.

To sum this section up, infection do not seem to affect network structure or complexity, neither in the case of bacteria or fungi populations. These results suggest that even though the microbe populations differ significantly between healthy and infected plants, the newly assembled communities are able to reach complexity levels compared to the controls.

## 4. Discussion

Studies examining how plant-associated microbiota respond to viral infections are still very scarce (Scortichini & Fiallo-Olivé, 2022). Our work shows that Turnip mosaic virus infection perturbs and restructures bacterial populations in the rhizosphere of *Arabidopsis thaliana*. Despite an overall reduction in bacterial diversity, microbial communities were able to withstand the stress imposed by viral infection, reestablishing interactions and, in some cases, increasing network complexity compared to control plants. The bacterial genera enriched under infection possess ecological traits allowing them to exploit this imbalance opportunistically, and in some cases, may indirectly support plant function. These results highlight the dynamic and resilient nature of the root microbiome under biotic stress, providing one of the few examples linking plant viral infection to microbial community re-assembly.

Host genotype further modulates these responses. It is well established that root and rhizosphere microbial communities vary with host genetic variation(S. Deng et al., 2021), and our findings support this pattern. In our Arabidopsis thaliana genotypes (Col⍰0 vs Mar⍰12), it is plausible that differences in root exudation profiles determined by genotype — alter the rhizosphere chemical environment and thus select for different microbial taxa, as root exudate variation is a known driver of microbiome assembly (Jacoby et al., 2017; Sasse et al., 2018; Zhalnina et al., 2018). Given that plant genotypes often differ in their tolerance and/or resistance to viral infection (Pagán & García-Arenal, 2018), the physiological responses to infection (including changes in exudation) are likely genotype⍰specific, leading to the recruitment of different microbial consortia under stress. Therefore, the taxa enriched upon infection in each genotype likely reflect a genotype-specific re⍰assembly of the microbiome triggered by the interaction between host genotype and stress.

One of the main effects of viral infection was a significant reduction in the diversity of bacterial communities, whereas fungal communities remained largely unaffected. Similarly, the population structure of bacteria diverged depending on plant health status, whereas fungal community structure remained stable. This pattern is consistent with previous findings showing that bacterial communities often respond more rapidly and flexibly than fungal communities to environmental disturbances. For example, bacterial communities changed much more than fungal communities along a steep wood⍰ash⍰driven pH gradient (Cruz-Paredes et al., 2021), and studies regarding water warming have observed stronger structural shifts in bacteria than fungi (X. Wu et al., 2024). The differential response likely reflects fundamental ecological and life⍰history differences: bacteria tend to exploit simple substrates quickly and proliferate, while fungi, with slower growth and different resource use, form more stable, specialized communities (C. Wang & Kuzyakov, 2024). Thus, the observed bottleneck and re⍰assembly of bacterial, but not fungal, diversity after infection is biologically plausible and aligns with broad patterns in microbial ecology.

Analysis of enriched bacterial genera further supports this interpretation. In Col-0 plants, the genera *Pelosinus, Actinotalea*, and *Rhizobium* were enriched under viral infection. The increase of *Pelosinus* and *Actinotalea* is most likely explained by their capacity to act as ecological opportunists. *Pelosinus* is known for thriving in disturbed or nutrient-poor environments and for its fermentative, stress-tolerant metabolism, which allows it to occupy niches that become available when microbial diversity declines (Mosher et al., 2012). Similarly, *Actinotalea*, a genus of oligotrophic *Actinobacteria*, often proliferates in soils with reduced carbon inputs or under environmental stress, reflecting its metabolic ability to grow efficiently under resource limitation (Semenova et al., 2022). In both cases, enrichment does not necessarily imply a functional role in plant–pathogen interactions, but rather the ability to exploit altered rhizosphere conditions generated by viral infection. In contrast, enrichment of Rhizobium may reflect a more directional response. Although some Rhizobium species are opportunistic root colonizers, several nonsymbiotic strains have demonstrated plant-growth-promoting activities, including hormone production and enhanced stress tolerance. Moreover, increases of Rhizobium in roots and endosphere have been reported under drought and other abiotic stress conditions, suggesting that this genus can proliferate when environmental conditions shift (Legeay et al., 2024). Thus, their increased abundance may indicate that the plant selectively benefits from, or at least tolerates, them under infection.

In the Mar-12 genotype, *Methylotenera* and *Flavobacterium* were enriched. Both genera possess metabolic traits that favour opportunistic expansion following stress-induced shifts in root exudation. *Methylotenera* is an obligate methylamineutilizing methylotroph (Kalyuzhnaya et al., 2006), and methylotrophic bacteria more broadly are frequently associated with plant surfaces and rhizospheres and have been implicated in plant stress responses (Knief et al., 2012; Kumar et al., 2019). Direct reports of *Methylotenera* increasing in plant rhizospheres under stress are limited, so here its enrichment should be interpreted as a plausible opportunistic response grounded in its C1-metabolism. *Flavobacterium*, by contrast, has well-documented abilities to degrade plant polysaccharides (e.g., pectin) and to expand its colonization in response to such substrates (Kraut-Cohen et al., 2021), and some strains have been shown experimentally to improve plant drought tolerance (Kim et al., 2023). Together, these lines of evidence support the interpretation that *Methylotenera* and *Flavobacterium* can exploit altered exudation or resource landscapes in stressed plants, with stronger experimental backing for stress-related plant benefits in the case of *Flavobacterium*. Thus, in Mar-12, the enrichment of these genera likely reflects a combination of opportunistic metabolic specialization and sensitivity to plant physiological stress signals, together shaping the restructured microbial community. The fact that *Haliangium* was enriched in both genotypes is especially noteworthy, it suggests a consistent ecological advantage under root⍰microbiome disruption. As a myxobacterium known to prey on other bacteria and to produce bioactive secondary metabolites (e.g., antifungal haliangicin), *Haliangium* could exploit empty niches and degraded microbiota, expanding opportunistically after rhizosphere perturbation (Fudou et al., 2002; Zhang et al., 2023).

As stated early, viral infection imposes a clear stress on the native rhizosphere bacterial community, likely creating a population bottleneck as susceptible taxa decline. Our network analyses indicate that, despite this perturbation, the surviving bacterial populations are capable of reorganizing and establishing new interactions, sometimes achieving network complexities comparable to or even higher than those observed in healthy plants. This pattern is consistent with findings in other plant–virus systems: for instance, Reovirus infection in rice triggers microbial reassembly via metabolite-mediated recruitment and exclusion, leading to novel community interactions (Li et al., 2025), while Sugarcane mosaic virus reduces bacterial diversity and network complexity in maize roots, highlighting the sensitivity of bacterial networks to viral stress (Liu et al., 2023). In our data, bacterial networks in both Col-0 and Mar-12 were dominated by *Alphaproteobacteria* and *Gammaproteobacteria*, as the enriched genera during infection frequently belonged to these classes. In contrast, fungal networks appeared largely unaffected, reflecting the relative stability of fungal populations and their lower responsiveness to rapid environmental or host-induced perturbations. Overall, these results suggest that viral infection acts as a selective filter on bacterial communities, yet the surviving taxa can reorganize into functional networks, maintaining or even enhancing connectivity despite the initial stress.

In summary, this study provides one of the few comprehensive characterizations of how a plant virus reshapes the rhizosphere microbiome. By demonstrating genotype-specific reassembly, opportunistic expansion of resilient bacterial taxa, and the ability of surviving communities to reorganize into complex networks, our findings advance understanding of microbiome dynamics under biotic stress and highlight mechanisms of microbial resilience and adaptability.

---

**Figure S1.**
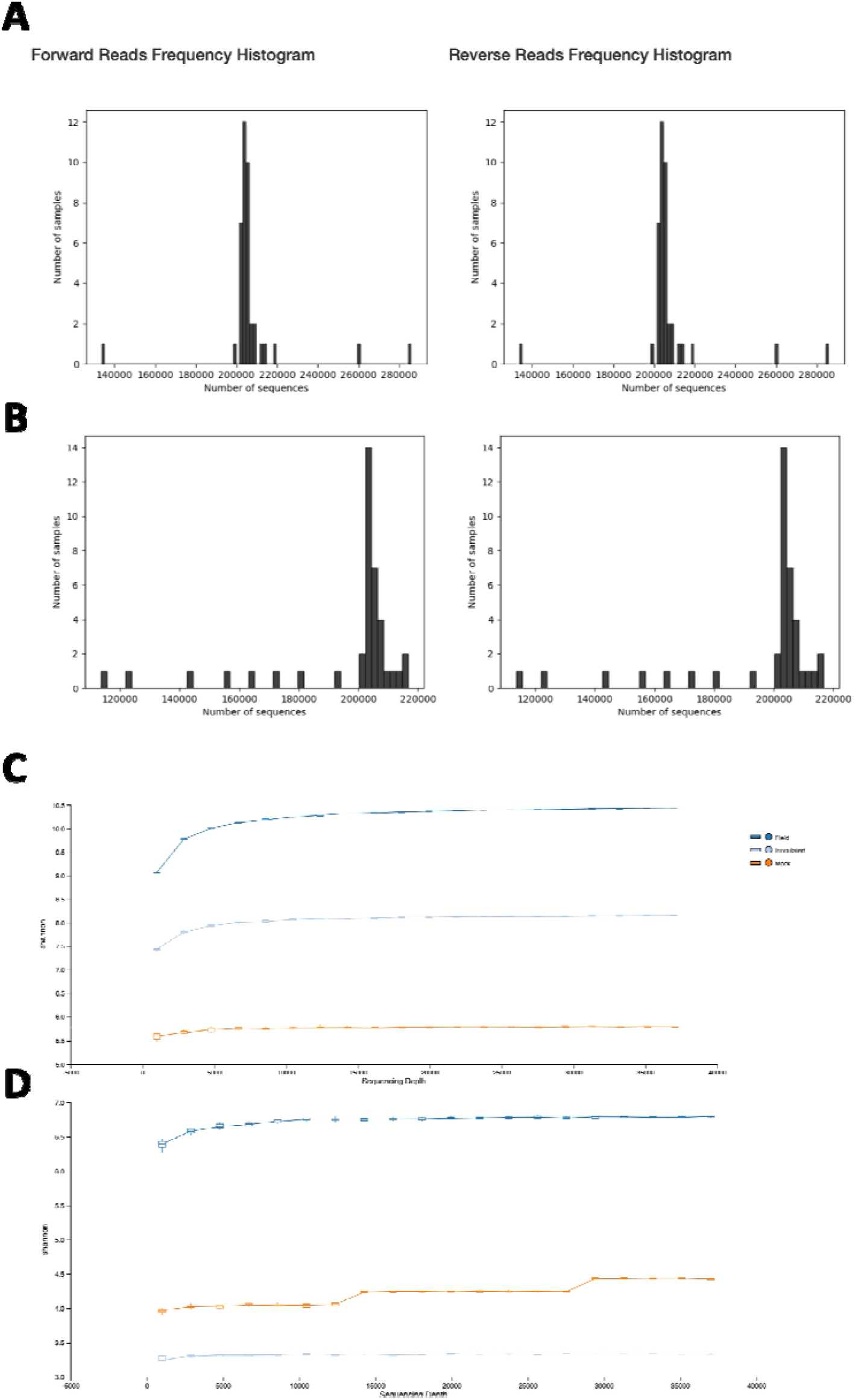
Description of the amplicon quality of our raw data. A) Read distribution across bacterial ASVs. B) Read distribution across fungal ASVs. C) Alpha rarefaction plot of the bacteria 16S sequencing. D) Alpha rarefaction of the fungi ITS-1 sequencing.

## Notes

### Competing Interest Statement

The authors have declared no competing interest.

### Summary of Updates

This revision was made to add one author to the manuscript.

### Funding

- MICIU/AEI/10.13039/501100011033 and European Union NextGenerationEU/PRTR: CNS2022-135350

